# SARS-CoV-2 ORF1ab^A1061S^ potentiate autoreactive T cell responses via epitope mimicry: an explanation to hepatitis of unknown cause

**DOI:** 10.1101/2022.05.16.491922

**Authors:** Yu Wang, Yuexing Liu

**Affiliations:** Shanghai Institute of Nutrition and Health, Shanghai Institutes for Biological Sciences, University of Chinese Academy of Sciences, Chinese Academy of Sciences; Shanghai 200031, China; Guangzhou Laboratory, Guangzhou 510005, China

**Keywords:** SARS-CoV-2, Hepatitis of unknown cause, Epitope mimicry, ORF1ab, A1061S mutation

## Abstract

The World Health Organization have recently announced outbreak news of acute, severe hepatitis of unknown cause in children under a Covid-19 pandemic. Whether it is associated with severe acute respiratory syndrome coronavirus 2 (SARS-CoV-2) infection is still under debating. Here, we performed genomic sequence alignment analysis of the genome of SARS-Cov-2 (Wuhan-hu-1) to the human genome reference. Sequence analysis revealed that the SARS-CoV-2 ORF1ab^1056-1173^ presented high identities with the human protein PAPR14^53-176^(3Q6Z_A). After searching the fully sequenced SARS-CoV-2 genomes deposited in GISAID (https://www.gisaid.org/), we detected 170 SARS-CoV-2 variants with mutation in ORF1ab^1061^, where alanine (A) was substituted by serine (S). This alteration made a 7-amino acid peptide (VVVNASN) in ORF1ab^1056-1062^ identical to its counterpart in PARP14^53-59^(3Q6Z_A). HLA prediction suggested that the peptides with high identities in PARP14 and ORF1ab could be presented by a same globally prevalent HLA-A*11:01 molecule. And in consistent with the first reported case of hepatitis of unknown, SARS-CoV-2 ORF1ab^VVVNASN^ variants were mostly identified as Delta lineages in UK by the late 2021, with an overall frequency of 0.00161%. Thus, our preliminary results raised a possibility that infection by SARS-CoV-2 ORF1ab^VVVNASN^ variant might elicit an autoimmune T cell response via epitope mimicry and is associated with the outbreak of unknown hepatitis. We anticipated that these findings will alert the human societies to pay more attention to rare mutations beyond the spike proteins.

## Main text

The severe acute respiratory syndrome coronavirus 2 (SARS-CoV-2) have caused a global pandemic in the last two years and are still evolving nowadays. Although we have made great progresses in understanding their virology and key steps of life cycle [1], long-term and systematic impacts of these virus on human are still missing, especially of that on the systemic immune status. Beside SARS-CoV-2, the World Health Organization have announced outbreak news of acute, severe hepatitis of unknown origin in children on 23, April 2022 [2]. And according to the records by European Centre for Disease Prevention and Control, the total number of cases reported worldwide were approximately 450 by 11, May 2022[3]. All of the patients were found in children under 16 years old, and it was estimated that more than 10% of the children have required liver transplantation [2].

The etiology of this disease is current unknown. However, some of these children have been identified with ongoing or recent SARS-CoV-2 infections in Israel and the USA [2]. Although linkes between long-lasting effects from the SARS-CoV-2 infection with hepatitis of unknown origin were currently without experimental examination, a case report highlighted a possible association between SARS-CoV-2 infection and subsequent development of T cell participated autoimmune liver disease[4]. In supporting, SARS-CoV-2 superantigens have been hypothesized to involve in excessive T cell activation and possibly the pathogenesis of hepatitis of unknown cause[5].

T cell responses are critically important to eliminate virus infection and also for SARS-CoV-2[6]. The elimination of virus finally resulted in a persistent T cell-pools that processing diverse T cell receptor (TCR) repertoires to virus related antigens, thus leaving a protection for the secondary infections by the same virus. Typically, these T cells were kept in check and did not attack human tissues. However, in some cases, virus infection led to immune disorders and caused autoimmune diseases, which were related to the disturbed peripheral tolerance[7, 8]. In the cases of SARS-CoV-2 infections, severe immune perturbations have been noted[9], thus rising possibilities that impaired peripheral T cell tolerance would occur.

Based on that knowledge, we hypothesized that virus mutations might generate new antigens that mimicked self-peptides and elicited an autoimmune response in appropriate inflammatory microenvironments, and thus associated with the emerging of hepatitis of unknown cause.

### PSI-blast identified a motif with high identity between human PARP14 and SARS-CoV-2 ORF1ab

To test the hypothesis above, we made a whole-genome alignment of SARS-CoV-2 (wh-hu-1)[10] to the human reference genome with PSI-blast in default parameters. Among all the SARS-CoV-2 proteins aligned, only a hit in ORF1ab (ORF1ab^1056-1173^) was found, which presented high identity with a sequence in human PARP14 (PARP14^53-176^, sequence ID: 3Q6Z_A) (**Fig. 1A**). The most identical peptide was located in a motif with 7 amino acids residuals, in which only a differential amino acid was found in ORF1ab^1061^ comparing to PARP14^58^ (**Fig. 1A**). In addition, PAPR14^62-75^(sequence ID: 3Q6Z_A) also shared high identity with ORF1ab^1065-1078^ (**Fig. 1A**).

**Figure 1.**
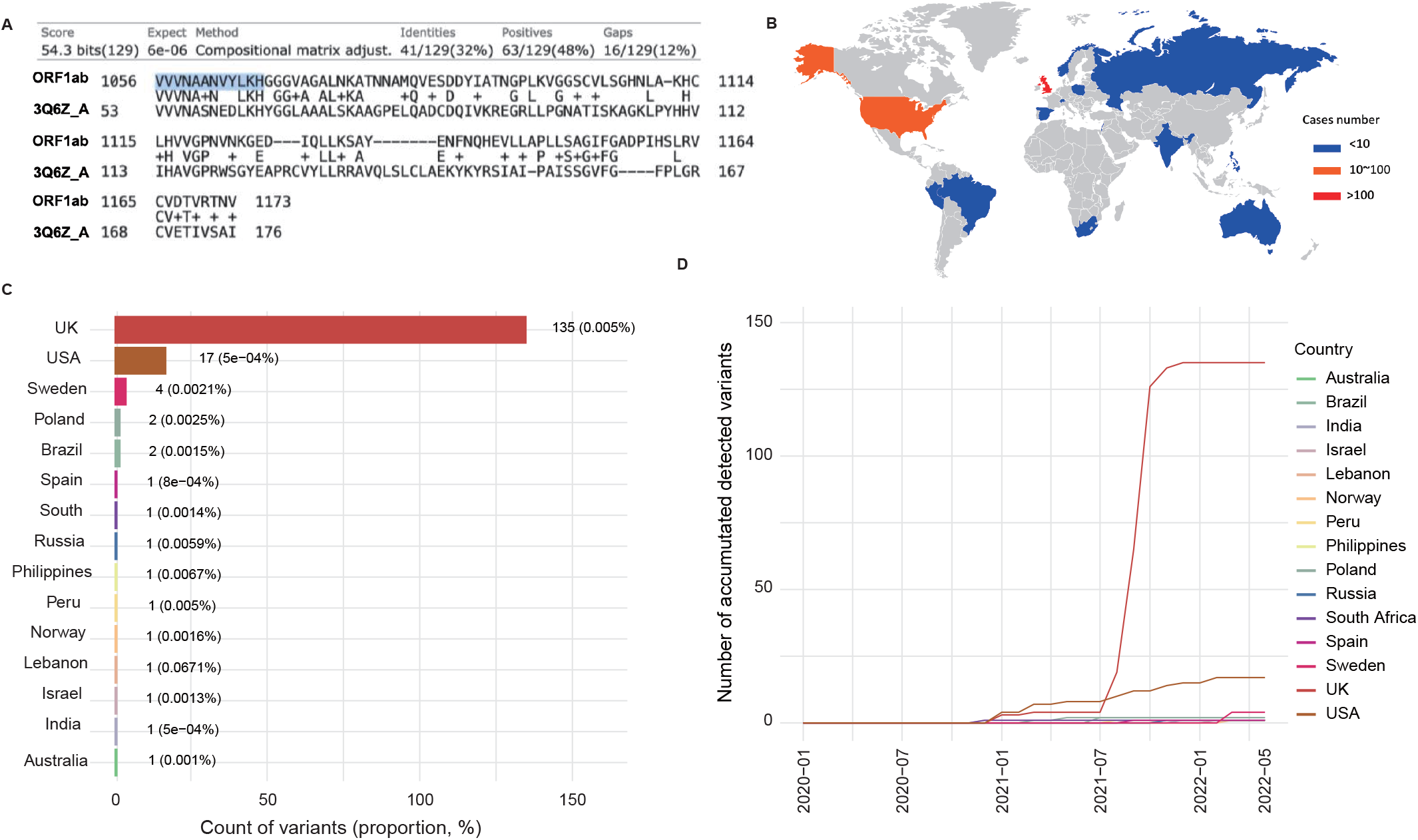
SARS-CoV-2 OFR1ab^A1061S^ substitution increased sequence identity with human PARP14 and variant’s epidemiology. **(A)** PSI blast showing the alignment of SARS-CoV-2 ORF1ab^1056-1173^ with PARP14^53-168^(3Q6Z_A). **(B)** Global distribution of the fully sequenced SARS-CoV-2 genomes possessing A1601S mutation in SARS-CoV-2 OFR1ab VVVNAAN motif. Basal layer map was created by BioRender.com with permission. **(C)** Counts (and proportions) of OFR1ab^VVVNASN^ variants across 15 countries. **(D)** Monthly trends in accumulated number of OFR1ab^VVVNASN^ variants across 15 countries.

### Rare mutation increased identity between ORF1ab^1056-1062^ and PARP14^53-59^

Since the release of the first sequence of SARS-CoV-2 (wuhan-hu-1) in December 2019, the virus has undergone numerous mutations. The accumulation of mutations in the genome of SARS-CoV-2 will affect functional properties and may alter infectivity, disease severity or interactions with host immunity. It was reasonable to speculate that there might be some SARS-CoV-2 variants with mutations occurring in ORF1ab^1056-1062^, thus resulting in an increased identity to human PARP14^53-59^ (3Q6Z_A). To test that hypothesis, we downloaded the proteins of SARS-CoV-2 variants deposited on GISAID (https://www.gisaid.org/). Sequence analysis was focus on the amino acid sequence from 1055 to 1077 on ORF1a protein. Among all the SARS-CoV-2 protein sequences (10,541,935 in total, by May 10^th^, 2022), 170 possessed an alanine (A) to serine (S) substitution on the site 1061 of ORF1ab (**Fig. 1B and 1C**). Noteworthily, this alteration made the core amino acid sequences identical to that of human PARP14^53-59^ (3Q6Z_A). Further analysis revealed that SARS-CoV-2 ORF1ab^VVVNASN^ variants has been recorded in 15 countries across 5 continents (**Fig. 1B and 1C**). The variants varied by countries with an overall frequency of 0.00161% worldwide (**Fig. 1C**). In consistent with the firstly reported cases of hepatitis of unknown, SARS-CoV-2 ORF1ab^VVVNASN^ variants was mostly detected in the UK (135) and the USA (18) (**Fig. 1D**).

In addition to ORF1ab^A1061S^ substitution, several other mutations that would potentially increase the sequence identity were also identified. For example, ORF1ab^G1073A^ have been emerging at a relative high frequency (**Fig. 2A, table 1**). This mutation led a 11-amino acid motif in ORF1ab (LKHGGGVAAAL) to be more similar in chemical properties, comparing to human PARP14 (LKHYGGLAAAL) (**Fig. 2A**). The 170 variants bearing A1061S mutation in ORF1ab were also annotated to the Variants of Concern (VOC) and most of mutational variants are found within SARS-CoV-2 Delta lineage (**Fig. 2B**). Although the Omicron variants were causing most of the infections globally, an Israel research group has warned that Delta variants were still undergoing circulating in parallel to Omicron variants and might maintain its circulation in future[11].

**Figure 2.**
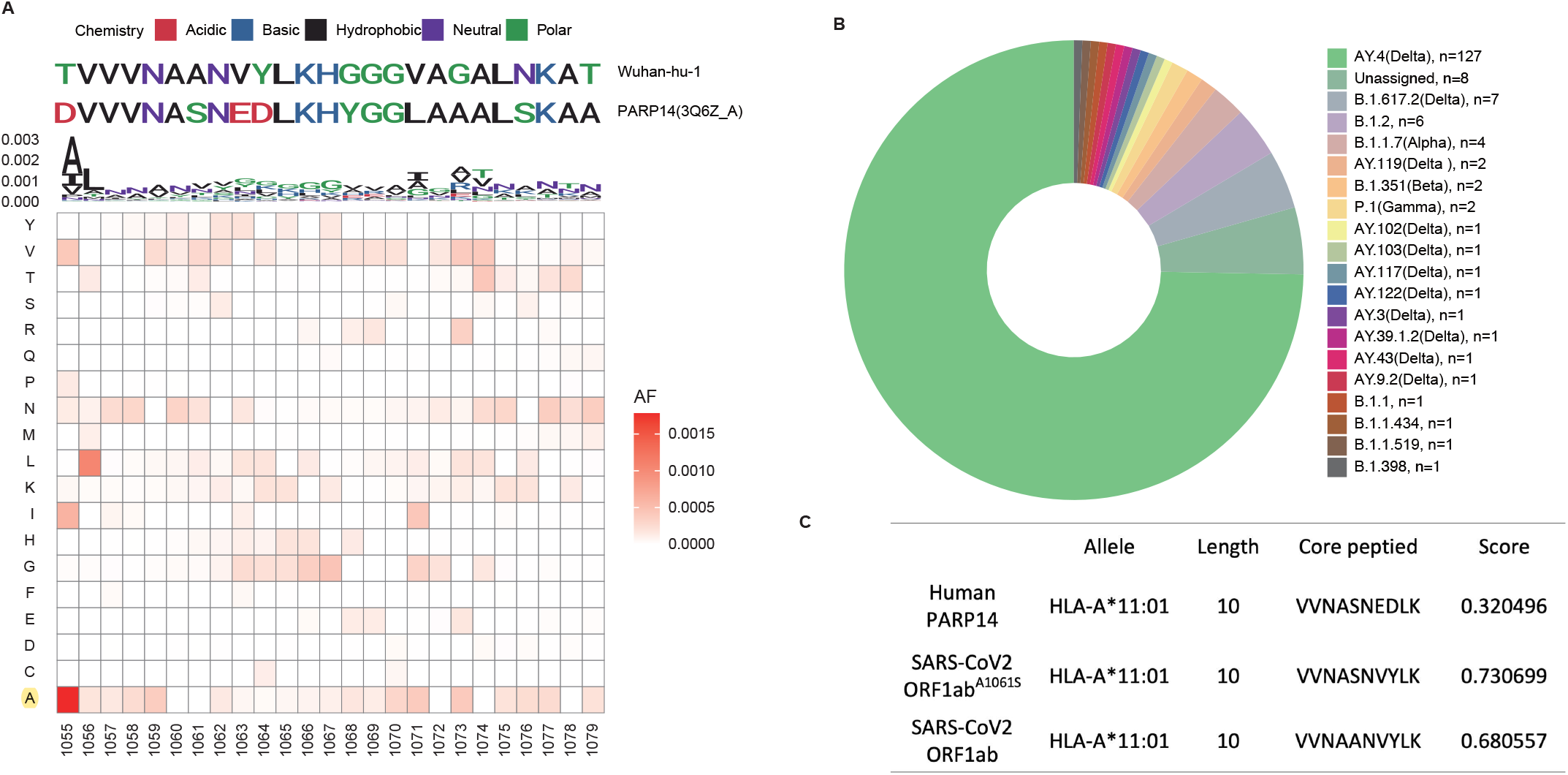
Sequence analysis of variants bearing OFR1ab^A1061S^ substitution. **(A)** Mutation preferences and rates in SARS-CoV-2 ORF1ab^1055-1079^ as compared to wuhan-hu-1. **(B)** Pie plot showing phylogenetic assignment of the SARS-CoV-2 variant (ORF1ab^VVVNASN^) with pangolin[15]. **(C)** Predicted binding ability by HLA-A*11:01 to human PARP14 and SARS-CoV-2 ORF1ab generated peptide. The MHCI binding predictions were made using the IEDB analysis resource NetMHCpan (ver. 4.1) tool [12].

**Table1.**
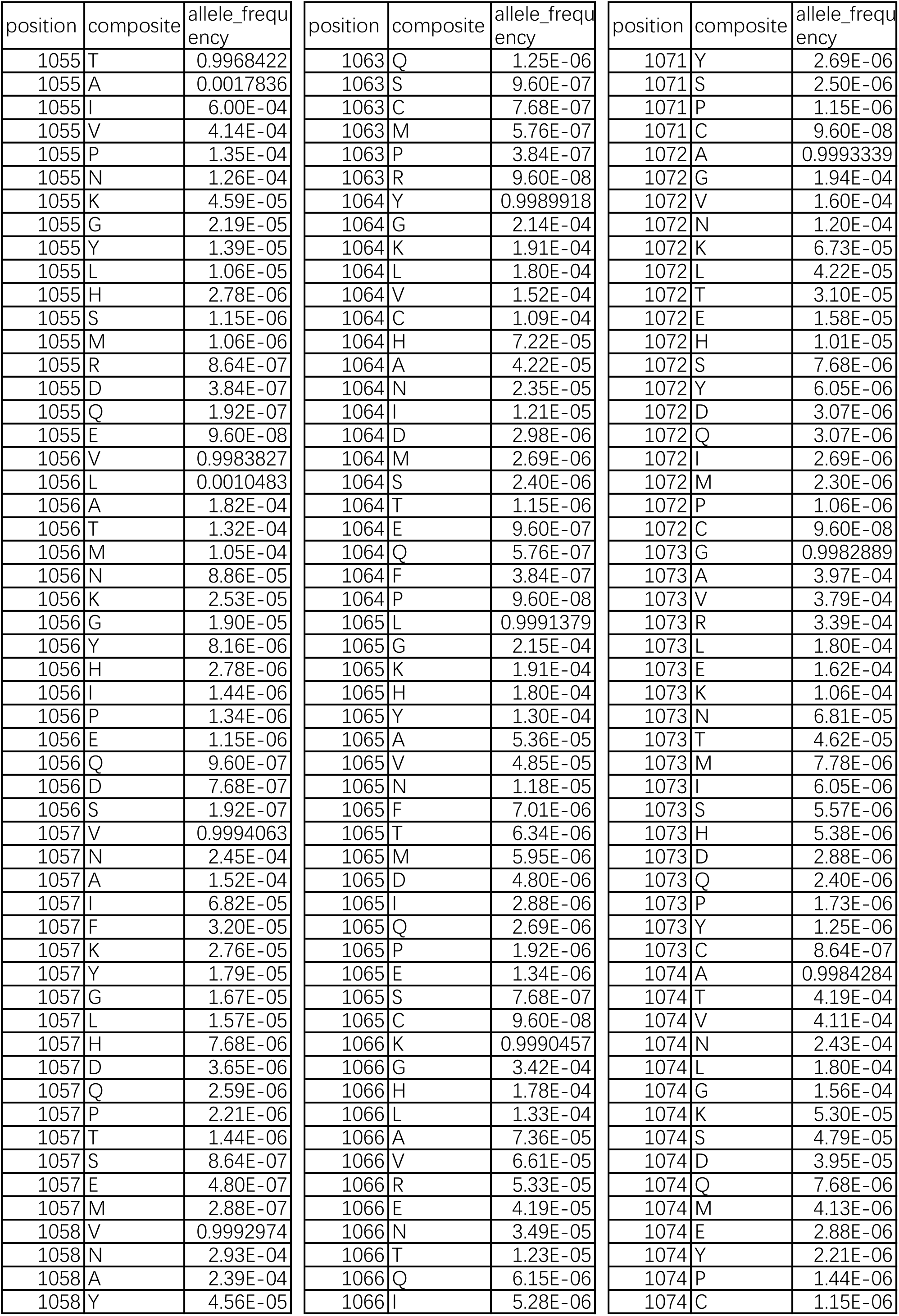
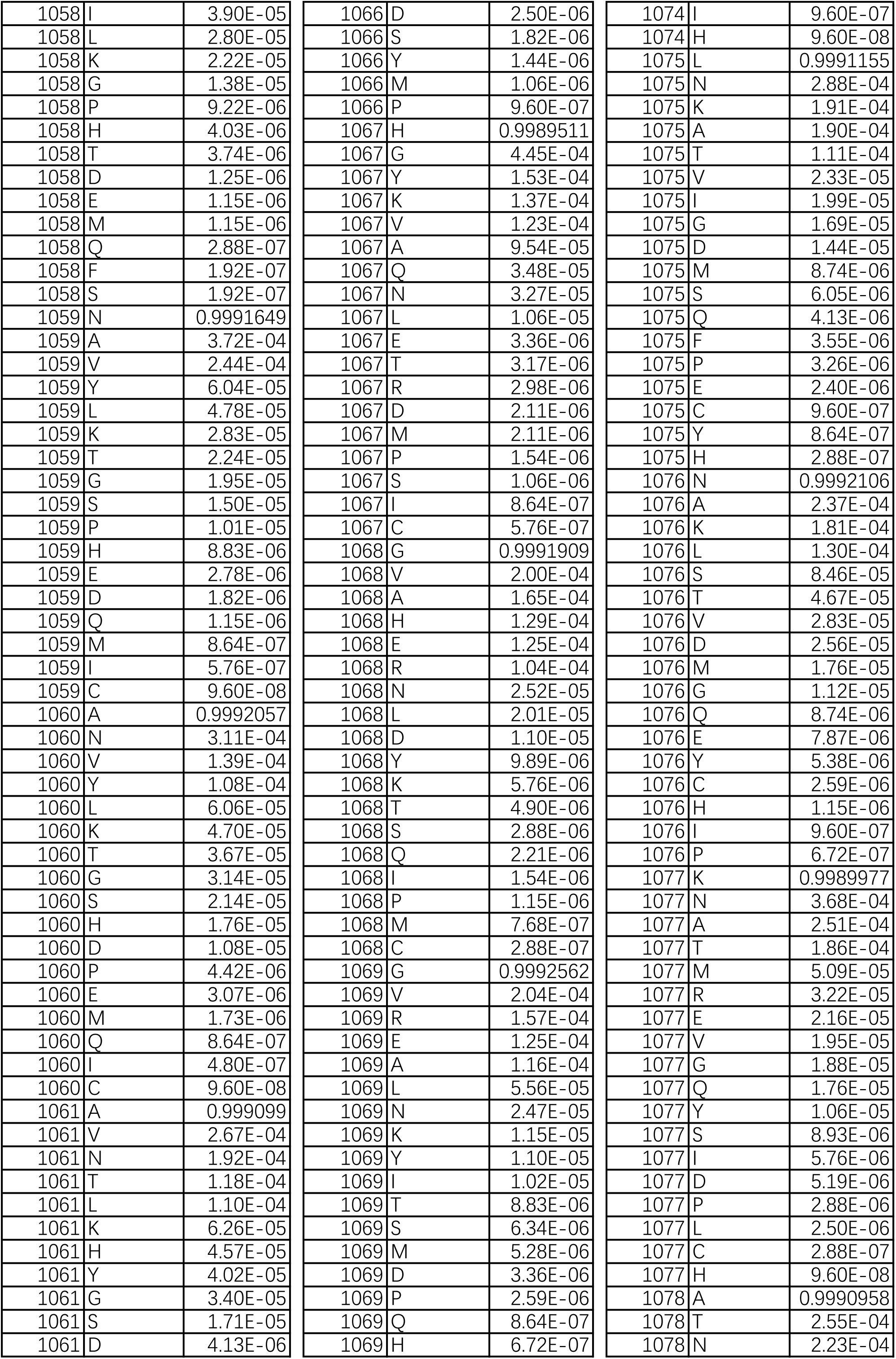
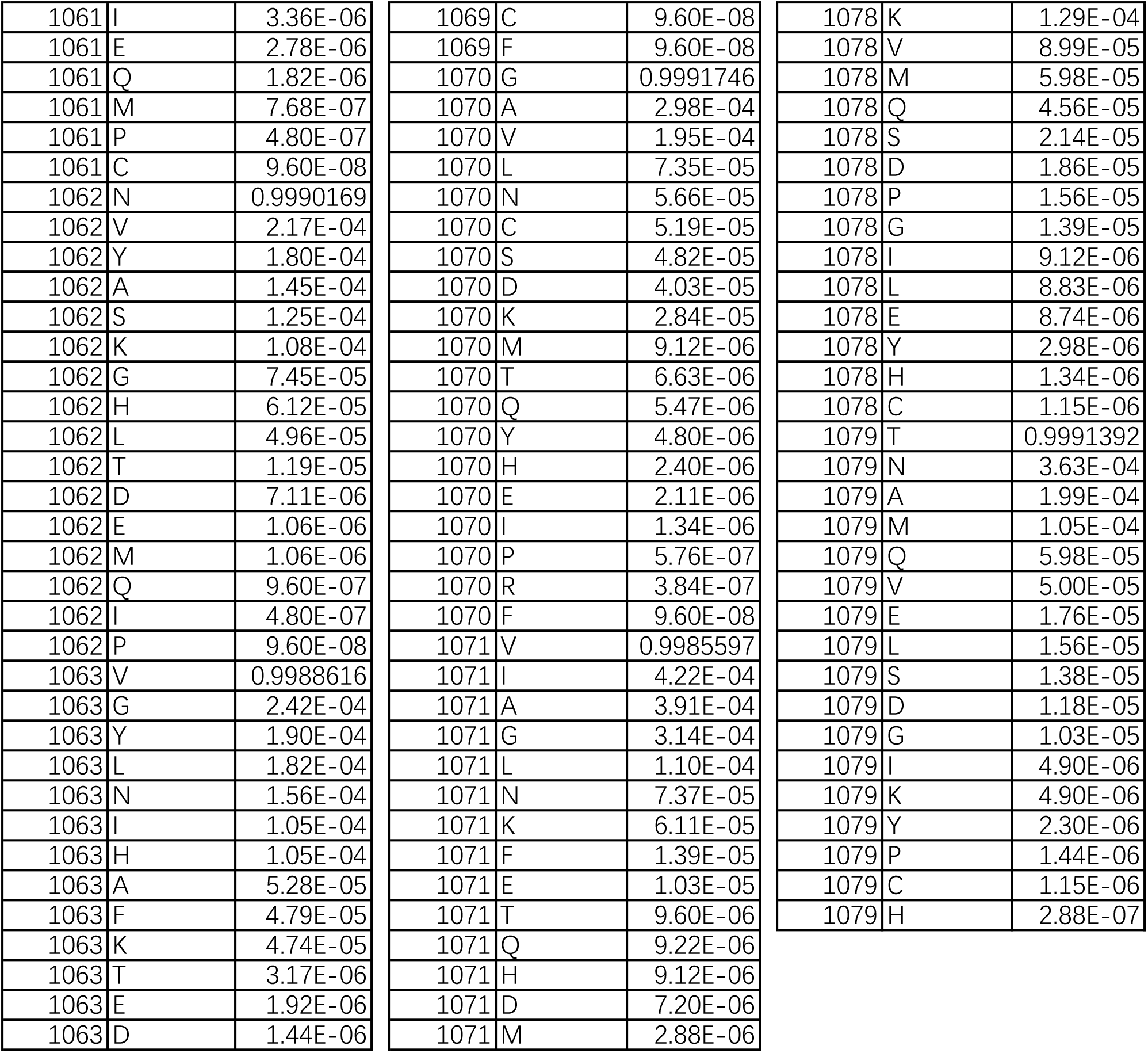
Mutation frequency in SARS-CoV-2 ORF1ab^1055-1079^ as compared to wuhan-hu-1.

| position | composite | allele_frequency | position | composite | allele_frequency | position | composite | allele_frequency |
| --- | --- | --- | --- | --- | --- | --- | --- | --- |
| 1055 | T | 0.9968422 | 1063 | Q | 1.25E-06 | 1071 | Y | 2.69E-06 |
| 1055 | A | 0.0017836 | 1063 | S | 9.60E-07 | 1071 | S | 2.50E-06 |
| 1055 | I | 6.00E-04 | 1063 | C | 7.68E-07 | 1071 | P | 1.15E-06 |
| 1055 | V | 4.14E-04 | 1063 | M | 5.76E-07 | 1071 | C | 9.60E-08 |
| 1055 | P | 1.35E-04 | 1063 | P | 3.84E-07 | 1072 | A | 0.9993339 |
| 1055 | N | 1.26E-04 | 1063 | R | 9.60E-08 | 1072 | G | 1.94E-04 |
| 1055 | K | 4.59E-05 | 1064 | Y | 0.9989918 | 1072 | V | 1.60E-04 |
| 1055 | G | 2.19E-05 | 1064 | G | 2.14E-04 | 1072 | N | 1.20E-04 |
| 1055 | Y | 1.39E-05 | 1064 | K | 1.91E-04 | 1072 | K | 6.73E-05 |
| 1055 | L | 1.06E-05 | 1064 | L | 1.80E-04 | 1072 | L | 4.22E-05 |
| 1055 | H | 2.78E-06 | 1064 | V | 1.52E-04 | 1072 | T | 3.10E-05 |
| 1055 | S | 1.15E-06 | 1064 | C | 1.09E-04 | 1072 | E | 1.58E-05 |
| 1055 | M | 1.06E-06 | 1064 | H | 7.22E-05 | 1072 | H | 1.01E-05 |
| 1055 | R | 8.64E-07 | 1064 | A | 4.22E-05 | 1072 | S | 7.68E-06 |
| 1055 | D | 3.84E-07 | 1064 | N | 2.35E-05 | 1072 | Y | 6.05E-06 |
| 1055 | Q | 1.92E-07 | 1064 | I | 1.21E-05 | 1072 | D | 3.07E-06 |
| 1055 | E | 9.60E-08 | 1064 | D | 2.98E-06 | 1072 | Q | 3.07E-06 |
| 1056 | V | 0.9983827 | 1064 | M | 2.69E-06 | 1072 | I | 2.69E-06 |
| 1056 | L | 0.0010483 | 1064 | S | 2.40E-06 | 1072 | M | 2.30E-06 |
| 1056 | A | 1.82E-04 | 1064 | T | 1.15E-06 | 1072 | P | 1.06E-06 |
| 1056 | T | 1.32E-04 | 1064 | E | 9.60E-07 | 1072 | C | 9.60E-08 |
| 1056 | M | 1.05E-04 | 1064 | Q | 5.76E-07 | 1073 | G | 0.9982889 |
| 1056 | N | 8.86E-05 | 1064 | F | 3.84E-07 | 1073 | A | 3.97E-04 |
| 1056 | K | 2.53E-05 | 1064 | P | 9.60E-08 | 1073 | V | 3.79E-04 |
| 1056 | G | 1.90E-05 | 1065 | L | 0.9991379 | 1073 | R | 3.39E-04 |
| 1056 | Y | 8.16E-06 | 1065 | G | 2.15E-04 | 1073 | L | 1.80E-04 |
| 1056 | H | 2.78E-06 | 1065 | K | 1.91E-04 | 1073 | E | 1.62E-04 |
| 1056 | I | 1.44E-06 | 1065 | H | 1.80E-04 | 1073 | K | 1.06E-04 |
| 1056 | P | 1.34E-06 | 1065 | Y | 1.30E-04 | 1073 | N | 6.81E-05 |
| 1056 | E | 1.15E-06 | 1065 | A | 5.36E-05 | 1073 | T | 4.62E-05 |
| 1056 | Q | 9.60E-07 | 1065 | V | 4.85E-05 | 1073 | M | 7.78E-06 |
| 1056 | D | 7.68E-07 | 1065 | N | 1.18E-05 | 1073 | I | 6.05E-06 |
| 1056 | S | 1.92E-07 | 1065 | F | 7.01E-06 | 1073 | S | 5.57E-06 |
| 1057 | V | 0.9994063 | 1065 | T | 6.34E-06 | 1073 | H | 5.38E-06 |
| 1057 | N | 2.45E-04 | 1065 | M | 5.95E-06 | 1073 | D | 2.88E-06 |
| 1057 | A | 1.52E-04 | 1065 | D | 4.80E-06 | 1073 | Q | 2.40E-06 |
| 1057 | I | 6.82E-05 | 1065 | I | 2.88E-06 | 1073 | P | 1.73E-06 |
| 1057 | F | 3.20E-05 | 1065 | Q | 2.69E-06 | 1073 | Y | 1.25E-06 |
| 1057 | K | 2.76E-05 | 1065 | P | 1.92E-06 | 1073 | C | 8.64E-07 |
| 1057 | Y | 1.79E-05 | 1065 | E | 1.34E-06 | 1074 | A | 0.9984284 |
| 1057 | G | 1.67E-05 | 1065 | S | 7.68E-07 | 1074 | T | 4.19E-04 |
| 1057 | L | 1.57E-05 | 1065 | C | 9.60E-08 | 1074 | V | 4.11E-04 |
| 1057 | H | 7.68E-06 | 1066 | K | 0.9990457 | 1074 | N | 2.43E-04 |
| 1057 | D | 3.65E-06 | 1066 | G | 3.42E-04 | 1074 | L | 1.80E-04 |
| 1057 | Q | 2.59E-06 | 1066 | H | 1.78E-04 | 1074 | G | 1.56E-04 |
| 1057 | P | 2.21E-06 | 1066 | L | 1.33E-04 | 1074 | K | 5.30E-05 |
| 1057 | T | 1.44E-06 | 1066 | A | 7.36E-05 | 1074 | S | 4.79E-05 |
| 1057 | S | 8.64E-07 | 1066 | V | 6.61E-05 | 1074 | D | 3.95E-05 |
| 1057 | E | 4.80E-07 | 1066 | R | 5.33E-05 | 1074 | Q | 7.68E-06 |
| 1057 | M | 2.88E-07 | 1066 | E | 4.19E-05 | 1074 | M | 4.13E-06 |
| 1058 | V | 0.9992974 | 1066 | N | 3.49E-05 | 1074 | E | 2.88E-06 |
| 1058 | N | 2.93E-04 | 1066 | T | 1.23E-05 | 1074 | Y | 2.21E-06 |
| 1058 | A | 2.39E-04 | 1066 | Q | 6.15E-06 | 1074 | P | 1.44E-06 |
| 1058 | Y | 4.56E-05 | 1066 | I | 5.28E-06 | 1074 | C | 1.15E-06 |

|  |  |  |
| --- | --- | --- |
| 1058 | I | 3.90E-05 |
| 1058 | L | 2.80E-05 |
| 1058 | K | 2.22E-05 |
| 1058 | G | 1.38E-05 |
| 1058 | P | 9.22E-06 |
| 1058 | H | 4.03E-06 |
| 1058 | T | 3.74E-06 |
| 1058 | D | 1.25E-06 |
| 1058 | E | 1.15E-06 |
| 1058 | M | 1.15E-06 |
| 1058 | Q | 2.88E-07 |
| 1058 | F | 1.92E-07 |
| 1058 | S | 1.92E-07 |
| 1059 | N | 0.9991649 |
| 1059 | A | 3.72E-04 |
| 1059 | V | 2.44E-04 |
| 1059 | Y | 6.04E-05 |
| 1059 | L | 4.78E-05 |
| 1059 | K | 2.83E-05 |
| 1059 | T | 2.24E-05 |
| 1059 | G | 1.95E-05 |
| 1059 | S | 1.50E-05 |
| 1059 | P | 1.01E-05 |
| 1059 | H | 8.83E-06 |
| 1059 | E | 2.78E-06 |
| 1059 | D | 1.82E-06 |
| 1059 | Q | 1.15E-06 |
| 1059 | M | 8.64E-07 |
| 1059 | I | 5.76E-07 |
| 1059 | C | 9.60E-08 |
| 1060 | A | 0.9992057 |
| 1060 | N | 3.11E-04 |
| 1060 | V | 1.39E-04 |
| 1060 | Y | 1.08E-04 |
| 1060 | L | 6.06E-05 |
| 1060 | K | 4.70E-05 |
| 1060 | T | 3.67E-05 |
| 1060 | G | 3.14E-05 |
| 1060 | S | 2.14E-05 |
| 1060 | H | 1.76E-05 |
| 1060 | D | 1.08E-05 |
| 1060 | P | 4.42E-06 |
| 1060 | E | 3.07E-06 |
| 1060 | M | 1.73E-06 |
| 1060 | Q | 8.64E-07 |
| 1060 | I | 4.80E-07 |
| 1060 | C | 9.60E-08 |
| 1061 | A | 0.999099 |
| 1061 | V | 2.67E-04 |
| 1061 | N | 1.92E-04 |
| 1061 | T | 1.18E-04 |
| 1061 | L | 1.10E-04 |
| 1061 | K | 6.26E-05 |
| 1061 | H | 4.57E-05 |
| 1061 | Y | 4.02E-05 |
| 1061 | G | 3.40E-05 |
| 1061 | S | 1.71E-05 |
| 1061 | D | 4.13E-06 |
| 1066 | D | 2.50E-06 |
| 1066 | S | 1.82E-06 |
| 1066 | Y | 1.44E-06 |
| 1066 | M | 1.06E-06 |
| 1066 | P | 9.60E-07 |
| 1067 | H | 0.9989511 |
| 1067 | G | 4.45E-04 |
| 1067 | Y | 1.53E-04 |
| 1067 | K | 1.37E-04 |
| 1067 | V | 1.23E-04 |
| 1067 | A | 9.54E-05 |
| 1067 | Q | 3.48E-05 |
| 1067 | N | 3.27E-05 |
| 1067 | L | 1.06E-05 |
| 1067 | E | 3.36E-06 |
| 1067 | T | 3.17E-06 |
| 1067 | R | 2.98E-06 |
| 1067 | D | 2.11E-06 |
| 1067 | M | 2.11E-06 |
| 1067 | P | 1.54E-06 |
| 1067 | S | 1.06E-06 |
| 1067 | I | 8.64E-07 |
| 1067 | C | 5.76E-07 |
| 1068 | G | 0.9991909 |
| 1068 | V | 2.00E-04 |
| 1068 | A | 1.65E-04 |
| 1068 | H | 1.29E-04 |
| 1068 | E | 1.25E-04 |
| 1068 | R | 1.04E-04 |
| 1068 | N | 2.52E-05 |
| 1068 | L | 2.01E-05 |
| 1068 | D | 1.10E-05 |
| 1068 | Y | 9.89E-06 |
| 1068 | K | 5.76E-06 |
| 1068 | T | 4.90E-06 |
| 1068 | S | 2.88E-06 |
| 1068 | Q | 2.21E-06 |
| 1068 | I | 1.54E-06 |
| 1068 | P | 1.15E-06 |
| 1068 | M | 7.68E-07 |
| 1068 | C | 2.88E-07 |
| 1069 | G | 0.9992562 |
| 1069 | V | 2.04E-04 |
| 1069 | R | 1.57E-04 |
| 1069 | E | 1.25E-04 |
| 1069 | A | 1.16E-04 |
| 1069 | L | 5.56E-05 |
| 1069 | N | 2.47E-05 |
| 1069 | K | 1.15E-05 |
| 1069 | Y | 1.10E-05 |
| 1069 | I | 1.02E-05 |
| 1069 | T | 8.83E-06 |
| 1069 | S | 6.34E-06 |
| 1069 | M | 5.28E-06 |
| 1069 | D | 3.36E-06 |
| 1069 | P | 2.59E-06 |
| 1069 | Q | 8.64E-07 |
| 1069 | H | 6.72E-07 |
| 1074 | I | 9.60E-07 |
| 1074 | H | 9.60E-08 |
| 1075 | L | 0.9991155 |
| 1075 | N | 2.88E-04 |
| 1075 | K | 1.91E-04 |
| 1075 | A | 1.90E-04 |
| 1075 | T | 1.11E-04 |
| 1075 | V | 2.33E-05 |
| 1075 | I | 1.99E-05 |
| 1075 | G | 1.69E-05 |
| 1075 | D | 1.44E-05 |
| 1075 | M | 8.74E-06 |
| 1075 | S | 6.05E-06 |
| 1075 | Q | 4.13E-06 |
| 1075 | F | 3.55E-06 |
| 1075 | P | 3.26E-06 |
| 1075 | E | 2.40E-06 |
| 1075 | C | 9.60E-07 |
| 1075 | Y | 8.64E-07 |
| 1075 | H | 2.88E-07 |
| 1076 | N | 0.9992106 |
| 1076 | A | 2.37E-04 |
| 1076 | K | 1.81E-04 |
| 1076 | L | 1.30E-04 |
| 1076 | S | 8.46E-05 |
| 1076 | T | 4.67E-05 |
| 1076 | V | 2.83E-05 |
| 1076 | D | 2.56E-05 |
| 1076 | M | 1.76E-05 |
| 1076 | G | 1.12E-05 |
| 1076 | Q | 8.74E-06 |
| 1076 | E | 7.87E-06 |
| 1076 | Y | 5.38E-06 |
| 1076 | C | 2.59E-06 |
| 1076 | H | 1.15E-06 |
| 1076 | I | 9.60E-07 |
| 1076 | P | 6.72E-07 |
| 1077 | K | 0.9989977 |
| 1077 | N | 3.68E-04 |
| 1077 | A | 2.51E-04 |
| 1077 | T | 1.86E-04 |
| 1077 | M | 5.09E-05 |
| 1077 | R | 3.22E-05 |
| 1077 | E | 2.16E-05 |
| 1077 | V | 1.95E-05 |
| 1077 | G | 1.88E-05 |
| 1077 | Q | 1.76E-05 |
| 1077 | Y | 1.06E-05 |
| 1077 | S | 8.93E-06 |
| 1077 | I | 5.76E-06 |
| 1077 | D | 5.19E-06 |
| 1077 | P | 2.88E-06 |
| 1077 | L | 2.50E-06 |
| 1077 | C | 2.88E-07 |
| 1077 | H | 9.60E-08 |
| 1078 | A | 0.9990958 |
| 1078 | T | 2.55E-04 |
| 1078 | N | 2.23E-04 |
| 1061 | I | 3.36E-06 |
| 1061 | E | 2.78E-06 |
| 1061 | Q | 1.82E-06 |
| 1061 | M | 7.68E-07 |
| 1061 | P | 4.80E-07 |
| 1061 | C | 9.60E-08 |
| 1062 | N | 0.9990169 |
| 1062 | V | 2.17E-04 |
| 1062 | Y | 1.80E-04 |
| 1062 | A | 1.45E-04 |
| 1062 | S | 1.25E-04 |
| 1062 | K | 1.08E-04 |
| 1062 | G | 7.45E-05 |
| 1062 | H | 6.12E-05 |
| 1062 | L | 4.96E-05 |
| 1062 | T | 1.19E-05 |
| 1062 | D | 7.11E-06 |
| 1062 | E | 1.06E-06 |
| 1062 | M | 1.06E-06 |
| 1062 | Q | 9.60E-07 |
| 1062 | I | 4.80E-07 |
| 1062 | P | 9.60E-08 |
| 1063 | V | 0.9988616 |
| 1063 | G | 2.42E-04 |
| 1063 | Y | 1.90E-04 |
| 1063 | L | 1.82E-04 |
| 1063 | N | 1.56E-04 |
| 1063 | I | 1.05E-04 |
| 1063 | H | 1.05E-04 |
| 1063 | A | 5.28E-05 |
| 1063 | F | 4.79E-05 |
| 1063 | K | 4.74E-05 |
| 1063 | T | 3.17E-06 |
| 1063 | E | 1.92E-06 |
| 1063 | D | 1.44E-06 |
| 1069 | C | 9.60E-08 |
| 1069 | F | 9.60E-08 |
| 1070 | G | 0.9991746 |
| 1070 | A | 2.98E-04 |
| 1070 | V | 1.95E-04 |
| 1070 | L | 7.35E-05 |
| 1070 | N | 5.66E-05 |
| 1070 | C | 5.19E-05 |
| 1070 | S | 4.82E-05 |
| 1070 | D | 4.03E-05 |
| 1070 | K | 2.84E-05 |
| 1070 | M | 9.12E-06 |
| 1070 | T | 6.63E-06 |
| 1070 | Q | 5.47E-06 |
| 1070 | Y | 4.80E-06 |
| 1070 | H | 2.40E-06 |
| 1070 | E | 2.11E-06 |
| 1070 | I | 1.34E-06 |
| 1070 | P | 5.76E-07 |
| 1070 | R | 3.84E-07 |
| 1070 | F | 9.60E-08 |
| 1071 | V | 0.9985597 |
| 1071 | I | 4.22E-04 |
| 1071 | A | 3.91E-04 |
| 1071 | G | 3.14E-04 |
| 1071 | L | 1.10E-04 |
| 1071 | N | 7.37E-05 |
| 1071 | K | 6.11E-05 |
| 1071 | F | 1.39E-05 |
| 1071 | E | 1.03E-05 |
| 1071 | T | 9.60E-06 |
| 1071 | Q | 9.22E-06 |
| 1071 | H | 9.12E-06 |
| 1071 | D | 7.20E-06 |
| 1071 | M | 2.88E-06 |
| 1078 | K | 1.29E-04 |
| 1078 | V | 8.99E-05 |
| 1078 | M | 5.98E-05 |
| 1078 | Q | 4.56E-05 |
| 1078 | S | 2.14E-05 |
| 1078 | D | 1.86E-05 |
| 1078 | P | 1.56E-05 |
| 1078 | G | 1.39E-05 |
| 1078 | I | 9.12E-06 |
| 1078 | L | 8.83E-06 |
| 1078 | E | 8.74E-06 |
| 1078 | Y | 2.98E-06 |
| 1078 | H | 1.34E-06 |
| 1078 | C | 1.15E-06 |
| 1079 | T | 0.9991392 |
| 1079 | N | 3.63E-04 |
| 1079 | A | 1.99E-04 |
| 1079 | M | 1.05E-04 |
| 1079 | Q | 5.98E-05 |
| 1079 | V | 5.00E-05 |
| 1079 | E | 1.76E-05 |
| 1079 | L | 1.56E-05 |
| 1079 | S | 1.38E-05 |
| 1079 | D | 1.18E-05 |
| 1079 | G | 1.03E-05 |
| 1079 | I | 4.90E-06 |
| 1079 | K | 4.90E-06 |
| 1079 | Y | 2.30E-06 |
| 1079 | P | 1.44E-06 |
| 1079 | C | 1.15E-06 |
| 1079 | H | 2.88E-07 |

### HLA prediction suggested potential overlaps of HLA in binding human and virus peptides

T cells recognized peptides presented by specific array of HLA molecules. Thus, HLA overlapping was a pre-determinant for cross T cell reactivities induced by epitope mimicry. We used an online HLA binding prediction tool[12] (http://tools.iedb.org/mhci/) to predict peptide binding abilities by a default array of MHC-I molecules, allowing to evaluate the binding potential and usage overlaps of HLA molecules by high similar peptides from human PARP14 and SARS-CoV2 ORF1ab. We selected a total of 23 amino acid surrounding the identical 7-amino acid of ORF1ab^A1061S^ and PARP14 as inputs respectively. As expected, we found that the peptide VVNASNELK in human PARP14 and VVNASNVYK in ORF1ab ^A1061S^ could be presented by a same HLA-A*11:01 molecule with comparable high-affinity (**Fig. 2C**). Importantly, OFR1ab^A1061S^ mutation showed increased binding ability of this peptide to HLA-A*11:01 molecule, as comparing to its wuhan-hu-1 counterpart (**Fig. 2C**). These findings suggested both the self- and virus-peptides with high sequence identity could be presented by a same HLA molecule, supporting that specific T-cell clones restricted to an HLA-A*11:01 molecules might be cross-activated by virus peptides mimicry and led to autoimmune responses. Indeed, HLA-A*11:01 was one of the most prevalent HLAs across the world [8], suggesting virus antigen presentation by HLA-A*11:01 were generally applicable to the worldwide.

## Discussion

Deregulated T cell responses are common triggers of various autoimmune diseases. Epitope mimicry of host proteins by pathogen is a common inducer in susceptible individuals to induce biased immune response versus tolerance, leading to tissue damage[13]. Here we found a mutation in SARS-CoV-2 ORF1ab may lead to increased identity between pathogen peptides with human proteins, providing a computational evidence for understanding the leading cause of SARS-CoV-2 associated autoimmune diseases, and also provided a new thought on the outbreaks of children hepatitis of unknown cause under a background of SARS-CoV-2 pandemic.

The outbreak of hepatitis of unknown cause was currently restricted to children under 16 years old. It should be noted that in children, their thymus output was kept in stack, T cell repertoires were continuously making, and peripheral tolerance was under establishing [14]. The disturbance of systematic and local immune microenvironment by SARS-CoV2 infection was likely to affect the establishment of normal T cell tolerances, thus possibly explained the preference of children to this type of hepatitis. Based on that, we anticipated that HLA genotyping may facilitate to uncovering the real cause of hepatitis.

According to the fully sequenced genomes deposited in GISAID, the frequency of SARS-CoV-2 ORF1ab^VVVNASN^ variants was around 0.0000161. Among 517,648,631 confirmed SARS-CoV-2 cases by 15, May 2022 (https://covid19.who.int/), it was roughly estimated that 8334 people, irrespective of children or adults, might be subjected to risks of autoimmune T cell responses and possibly the hepatitis of unknown cause. However, other unknown factors may also participate to exacerbate disease morbidity and severity, and indeed we have noted some mutations will increase the sequence identity between ORF1ab^1065-1078^ and PAPR14^62-75^(3Q6Z_A). Most of the variants bearing A1061S substitution were in Delta lineage, and it should be noteworthy that Delta variants were still circulating in parallel with Omicron variants[11]. Thus, mutations in these sites should call for our great concerns. However, our results were still preliminary and we only aimed to discussing a possible association of SARS-CoV-2 infection with children acute hepatitis of unknown cause. Further experimental validation of this hypothesis presented here was urgently needed to figure out the nature of hepatitis of unknown cause.

## Acknowledgement

We thank GISAID for collecting and sharing the sequences of SARS-CoV-2 globally.

## Author contributions

Both Yu Wang and Yuexing Liu contributed to the conceptional design and data processing.

## Fundings and ethics

This manuscript was not funded by any sponsors. All of the data was generated from public available database and did not required ethics committee approval.

